# Microbial helpers allow cyanobacteria to thrive in ferruginous waters

**DOI:** 10.1101/2020.05.08.085001

**Authors:** Nadia Szeinbaum, Yael Toporek, Christopher T. Reinhard, Jennifer B. Glass

## Abstract

The Great Oxidation Event (GOE) was a rapid accumulation of oxygen in the atmosphere as a result of the photosynthetic activity of cyanobacteria. This accumulation reflected the pervasiveness of O_2_ on the planet’s surface, indicating that cyanobacteria had become ecologically successful in Archean oceans. Micromolar concentrations of Fe^2+^ in Archean oceans would have reacted with hydrogen peroxide, a byproduct of oxygenic photosynthesis, to produce hydroxyl radicals, which cause cellular damage. Yet cyanobacteria colonized Archean oceans extensively enough to oxygenate the atmosphere, which likely required protection mechanisms against the negative impacts of hydroxyl radical production in Fe^2+^-rich seas. We identify several factors that could have acted to protect early cyanobacteria from the impacts of hydroxyl radical production and hypothesize that microbial cooperation may have played an important role in protecting cyanobacteria from Fe^2+^ toxicity before the GOE. We found that several strains of facultative anaerobic heterotrophic bacteria (*Shewanella*) with ROS defense mechanisms increase the fitness of cyanobacteria (*Synechococcus*) in ferruginous waters. *Shewanella* species with manganese transporters provided the most protection. Our results suggest that a tightly regulated response to prevent Fe^2+^ toxicity could have been important for the colonization of ancient ferruginous oceans, particularly in the presence of high manganese concentrations, and may expand the upper bound for tolerable Fe^2+^ concentrations for cyanobacteria.

## Introduction

Earth’s first biogeochemical cycles were driven by anaerobic microorganisms (Canfield et al., 2006; Martin et al., 2018). At around 2.3 Ga, the Great Oxidation Event (GOE) resulted in the initial oxygenation of the atmosphere and surficial biosphere, which ultimately led to the modern dominance of aerobic organisms on Earth’s surface (Bar-On et al., 2018; Luo et al., 2016). Although biological O_2_ production was a prerequisite for the GOE (Haqq-Misra et al., 2011; Holland, 2002), oxygenic photosynthesis may have emerged in cyanobacteria hundreds of millions of years prior to the initial accumulation of O_2_ in Earth’s atmosphere (Cardona et al., 2019; Lalonde and Konhauser, 2015; Ossa Ossa et al., 2018; Planavsky et al., 2014). The delay between the emergence of cyanobacterial O_2_ production and O_2_ accumulation in the atmosphere may have been modulated by geophysical drivers (Catling et al., 2001; Holland, 2009; Lee et al., 2016), but may also reflect the time required for metabolic innovations to appear in early cyanobacteria or for the emergence of ecological linkages with other microbes facilitating the success of cyanobacteria (Blank and Sanchez-Baracaldo, 2010; Johnston et al., 2009; Lyons et al., 2014; Ozaki et al., 2019). Understanding how cyanobacteria cooperated with other microbes to colonize the Earth’s surface is thus essential to understand the ecology and tempo of the GOE.

The emergence of oxygenic photosynthesis in cyanobacteria occurred in the Archean (Chisholm, 2017; Kendall et al., 2010; Konhauser et al., 2011; Lalonde and Konhauser, 2015; Lyons et al., 2014; Olson et al., 2013; Planavsky et al., 2014; Reinhard et al., 2013b). The metabolic expansion of cyanobacteria before the GOE may reflect their transition from land to Fe^2+^-rich Archean oceans (Herrmann and Gehringer, 2019). This transition would have been physiologically challenging due to Fe^2+^ toxicity from its reactions with reactive oxygen species (ROS) produced during photosynthesis (Swanner et al., 2015a). Archean oceans likely contained tens to hundreds of micromolar Fe^2+^ within the ocean interior (Canfield, 2005; Derry, 2015; Drever, 1974; Holland, 1973; Song et al., 2017; Thompson et al., 2019), which would have reacted rapidly in the surface ocean with O_2_ produced from photosynthesis and any hydrogen peroxide (H_2_O_2_) from photochemical reactions between O_2_ and dissolved organic matter, as well as enzymes like superoxide dismutase (Hansel and Diaz, 2020; Zinser, 2018b). This Fe^2+^-driven reaction, known as the Fenton reaction, produces hydroxyl radicals (·OH; **Eq. 1**):

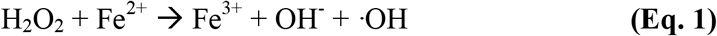

Hydroxyl radicals cause cellular damage, especially to DNA (Imlay, 2003; Imlay, 2008). Yet, cyanobacteria must have colonized vast areas of the ocean in order to oxygenate the atmosphere. Cyanobacteria may have been protected by spatial separation of oxygenic and anoxygenic phototrophs that could have removed upwelled ferrous iron prior to its arrival at the surface (Ozaki et al., 2019), though the potential effectiveness of this would depend on the Fe/P ratio of deep waters. Anti-oxidants such as elevated dissolved manganese (Mn^2+^) and ancient Mn-based catalases may have protected ancient cyanobacteria against ROS toxicity (Fischer et al., 2016; Lingappa et al., 2019).

Here, we test the hypothesis that heterotrophic microbial “helpers” may have protected cyanobacteria from ROS produced by Fenton chemistry in Archean oceans, thereby increasing cyanobacterial fitness and enabling their ecological success. Such microbial cooperation is common among modern cyanobacteria and heterotrophic proteobacteria (Christie-Oleza et al., 2017; Morris et al., 2011; Morris et al., 2008; Zinser, 2018a), whose intimate relationship is evidenced by extensive horizontal gene transfer (Ben Said and Or, 2017; Braakman et al., 2017; Goldenfeld and Woese, 2011). At the time of the GOE, many bacterial lineages, including Proteobacteria, had already diversified (Battistuzzi et al., 2004; Cavalier-Smith, 2006a; Cavalier-Smith, 2006b), which would have increased the phenotypic pool available for cooperation. Including microbial cooperation as an ecological mechanism in models of early Earth’s ecological history might provide a more realistic picture of the ancient interactions that ultimately led to the GOE.

We explored whether the presence of “helper” heterotrophic proteobacteria leads to increased fitness of cyanobacteria in ferruginous conditions. For a model cyanobacterium, we chose *Synechococcus* sp. PCC 7002 (hereafter *Synechococcus*), which was previously shown to experience Fe^2+^ toxicity at >100 µM Fe^2+^ associated with increased intracellular ROS production (Swanner et al., 2015a). As potential “helper” bacteria, we chose *Shewanella*, facultative anaerobic gammaproteobacteria that can survive O_2_ intrusions in the presence of high Fe^2+^ using diverse H_2_O_2_-scavenging enzymes (Jiang et al., 2014; Mishra and Imlay, 2012; Sekar et al., 2016). Experimental conditions loosely simulated a pre-GOE illuminated ferruginous surface ocean overlain by a CO_2_-and H_2_-rich anoxic atmosphere. We found that several *Shewanella* species allowed *Synechococcus* to grow in ferruginous conditions that significantly impaired growth of *Synechococcus* monocultures. The “helper” *Shewanella* strains all contained the ability to actively uptake dissolved manganese (Mn^2+^) via the natural resistance-associated macrophage protein (NRAMP) family MntH Mn^2+^ transporter, a strategy that has previously been shown to correlate with ROS survival (Daly et al., 2004). Our results stress the importance of considering microbial cooperation and alternative ROS strategies, such as manganese protection, in models of early Earth microbial ecology.

## Results

### Cyanobacteria growth is impaired in ferruginous conditions and is restored in the presence of some proteobacteria

We found that *Synechococcus* growth in the presence of elevated Fe^2+^ improved (to a varying degree) in the presence of all *Shewanella* spp. tested. In monoculture, *Synechococcus* had similar growth rate and yield at 25 and 500 μM Fe^2+^, but a longer lag period at 500 μM Fe^2+^ (∼2 days) than at 25 μM Fe^2+^ (∼1 day; **Fig. 1A)**. At 1000 μM Fe^2+^, *Synechococcus* growth was significantly impaired in monoculture, reaching only 10% the cell density of cultures with 25 and 500 μM Fe^2+^ **(Fig. 1A)**. *Synechococcus* growth was mostly recovered in the presence of high Fe^2+^ when grown in co-culture with *Shewanella baltica* OS-155, although the initial lag period was extended **(Fig. 1B)**. In the presence of *Shewanella algae* MN-01 **(Fig. 1C)** and *Shewanella loihica* PV-4 **(Fig. 1D)**, *Synechococcus* growth was partially recovered at high Fe^2+^. Other than an extended lag phase, *Shewanella algae* BrY **(Fig. 1E)** and *Shewanella oneidensis* MR-1 **(Fig. 1F)** had minimal influence on *Synechococcus* growth, compared to the monoculture **(Fig. 1A)**, in all three Fe^2+^ treatments. Although difficult to quantify due to spectral interference of Fe(III) oxide particles, *Shewanella* cell numbers declined throughout the experiment (data not shown).

**Figure 1.**
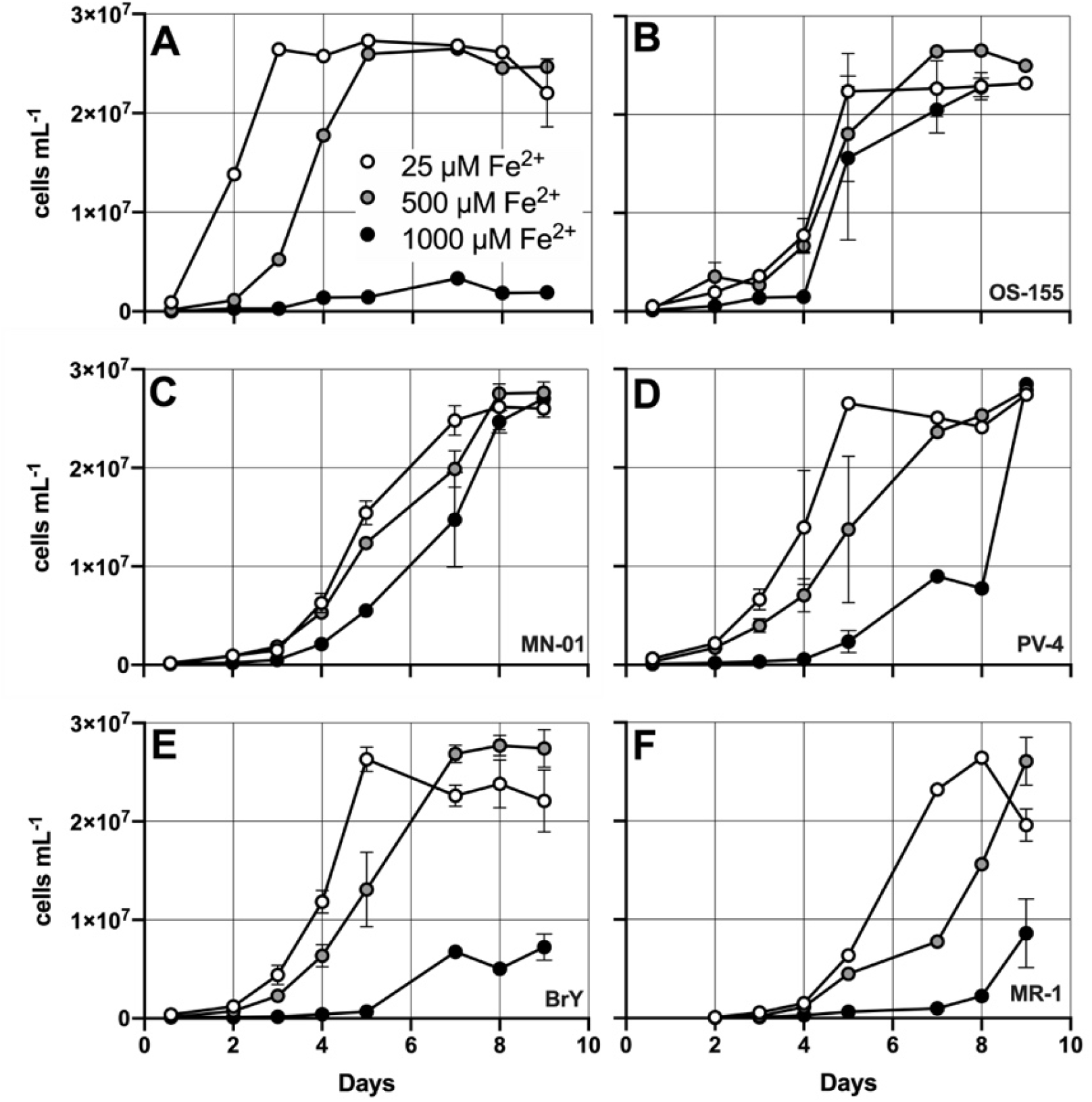
Growth of *Synechococcus* PCC 7002 in mono- or co-culture with *Shewanella* spp. with varying Fe^2+^. Co-cultures are: A) none, B) *Shewanella baltica* OS-155, C) *Shewanella algae* MN-01, D) *Shewanella loihica* PV-4, E) *Shewanella algae* BrY, F) *Shewanella oneidensis* MR-1. Error bars represent the standard error of the mean (n=3).

### The best proteobacterial helpers are the least H_2_O_2_ sensitive, and the best H_2_O_2_ scavengers

We measured growth and H_2_O_2_ scavenging rates of *Shewanella* spp. in the presence of varying H_2_O_2_. *S. baltica* OS-155 was the least sensitive to H_2_O_2_ **(Fig. 2A)**. *S. algae* MN-01 **(Fig. 2B)**, *S. loihica* PV-4 **(Fig. 2C)**, and *S. algae* BrY **(Fig. 2D)** were moderately sensitive to H_2_O_2_. *S. oneidensis* MR-01 was the most sensitive to H_2_O_2_ (**Fig. 2D)**. Along with being most H_2_O_2_ tolerant, *S. algae* MN-01 and *S. baltica* OS-155 had the highest rates of H_2_O_2_ scavenging activity, followed by *S. algae* BrY **(Fig. 3)**. *S. loihica* PV-4 and *S. oneidensis* MR-1 had the lowest H_2_O_2_ scavenging rates **(Fig. 3)**.

**Figure 2.**
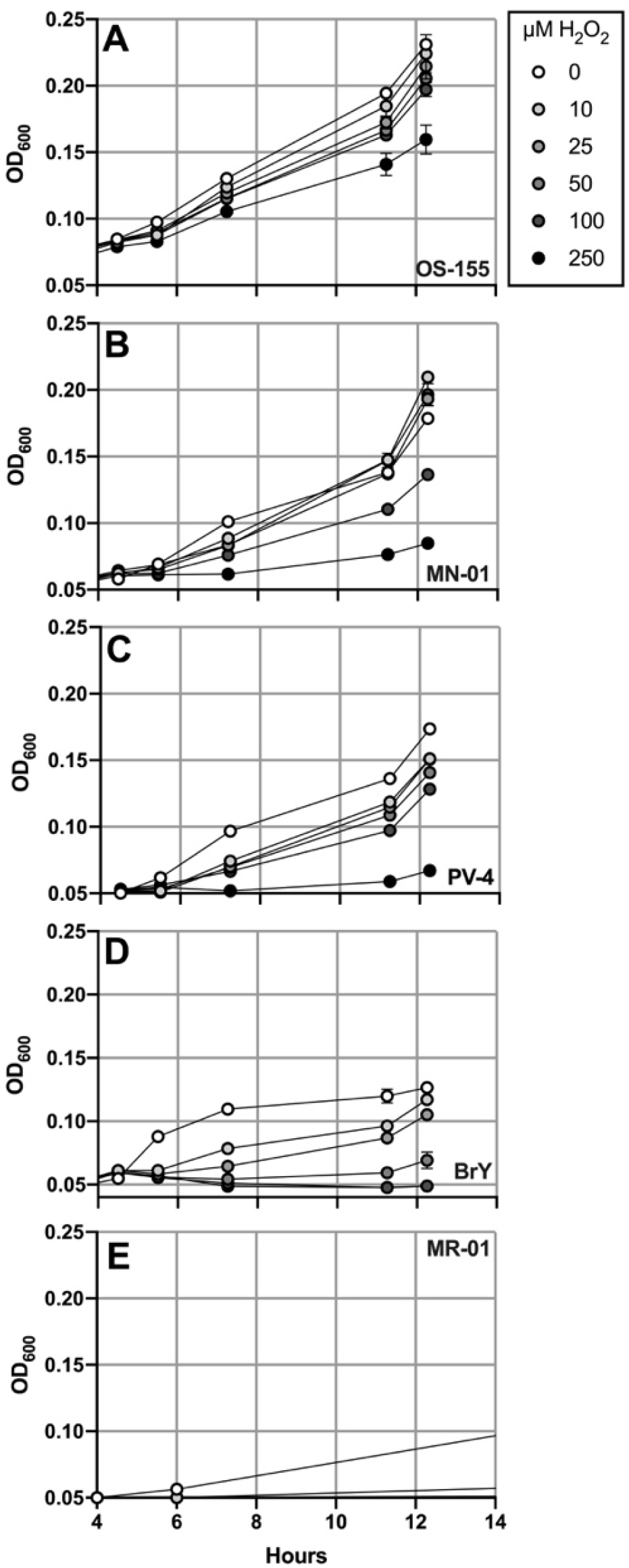
Growth of *Shewanella* spp. with varying H_2_O_2_. A) *Shewanella baltica* OS-155, B) *Shewanella algae* MN-01, C) *Shewanella loihica* PV-4, D) *Shewanella algae* BrY, E) *Shewanella oneidensis* MR-1. Error bars represent the standard error of the mean (n=3 for all except *S. baltica* OS-155, n=2). H_2_O_2_ was added at the four-hour timepoint.

**Figure 3.**
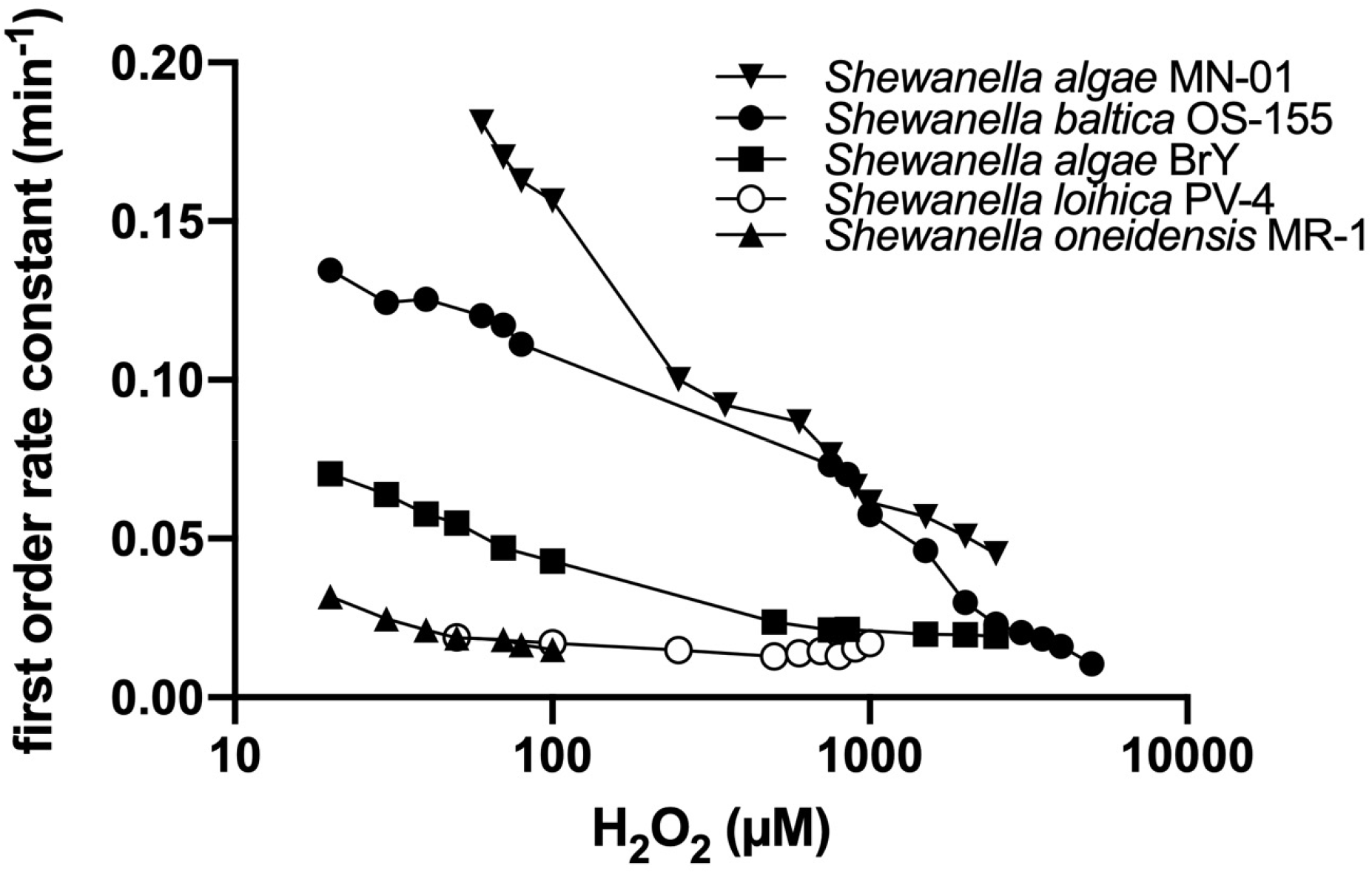
H_2_O_2_ peroxide scavenging capacity of *Shewanella* spp. shown as the first order rate constant plotted versus initial H_2_O_2_ concentration. No change in H_2_O_2_ was observed in the abiotic control.

### Manganese may protect cyanobacteria from Fe^2+^ toxicity

To test whether Mn^2+^ can protect cyanobacteria from Fe^2+^ toxicity, we grew *Synechococcus* PCC 7002 (four replicates per treatment) under anoxic conditions with the addition of 1 mM Fe^2+^ and/or 1 mM Mn^2+^. Cells with 1 mM Mn^2+^ grew similarly to the controls **(Fig. 4)**. The A+ medium contained background concentrations of ∼140 μM Fe^2+^ and ∼220 μM Mn^2+^. No growth occurred with 1 mM Fe^2+^. Red Fe(III) oxide particles indicated that Fe^2+^ had been oxidized and precipitated, as observed by Swanner et al. (2015b). Treatments with 1 mM Fe^2+^ and 1 mM Mn^2+^ resembled 1 mM Fe^2+^ treatments for approximately the first week. Between 4-13 days, two out of four of the Fe^2+^ and Mn^2+^ treatments grew to maximal OD_750_. In the next three months, another of the Fe^2+^ and Mn^2+^ treatments also turned green (data not shown). These results show that 1 mM Mn^2+^ is not toxic to cyanobacteria and may in fact aid in the survival of cyanobacteria Fe^2+^ toxicity, after an acclimation period. The mechanism underlying the apparent protective effect of Mn^2+^ that rescued growth for three out of four cultures in high Fe^2+^ after an extended lag phase remains unknown.

**Figure 4.**
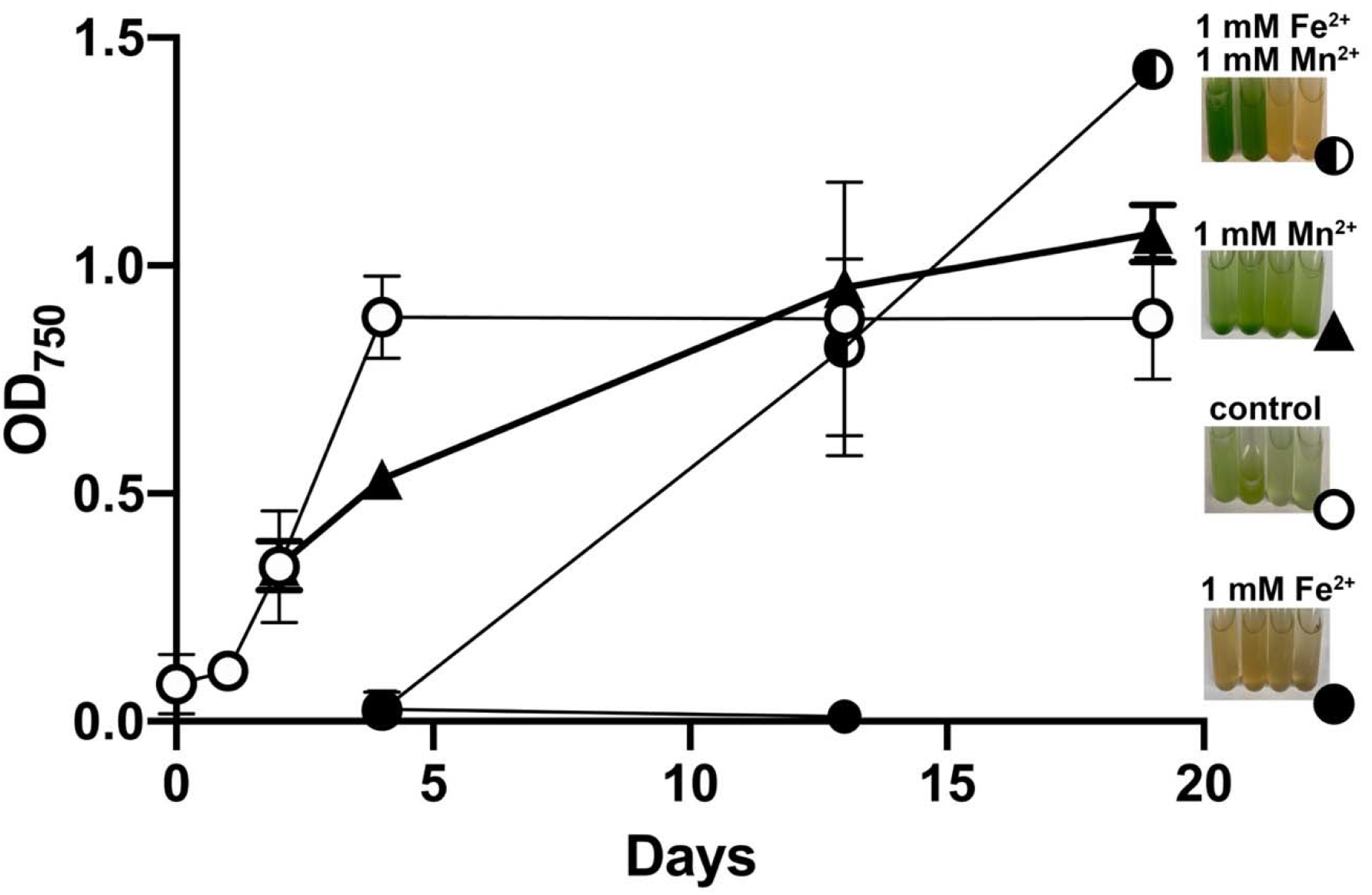
Growth of *Synechococcus* sp. PCC 7002 in monoculture with 1 mM Fe^2+^ and/or 1 mM Mn^2+^. Controls had background levels of ∼140 μM Fe^2+^ and ∼220 μM Mn^2+^. The growth curve for the 1 mM Fe^2+^ + 1 mM Mn^2+^ treatment is shown for the two replicates (out of four) that grew. Photos were taken on day 13.

### The best proteobacterial helpers encode additional genes for H_2_O_2_ degradation

*Synechococcus* PCC 7002’s susceptibility to Fe^2+^ toxicity is consistent with the limited number of catalase genes in its genome; it encodes cytoplasmic KatG but not periplasmic KatE **(Table 1)**. Without KatE to scavenge H_2_O_2_ in the periplasm, H_2_O_2_ can react with Fe^2+^ to generate ·OH intracellularly **(Eq. 1)**. Like *Synechococcus* PCC 7002, most marine cyanobacteria are KatE-negative; a BLAST search of cyanobacterial genomes in NCBI recovered KatE catalase homologs almost exclusively in freshwater and soil cyanobacteria (**Table S1)**.

**Table 1.**
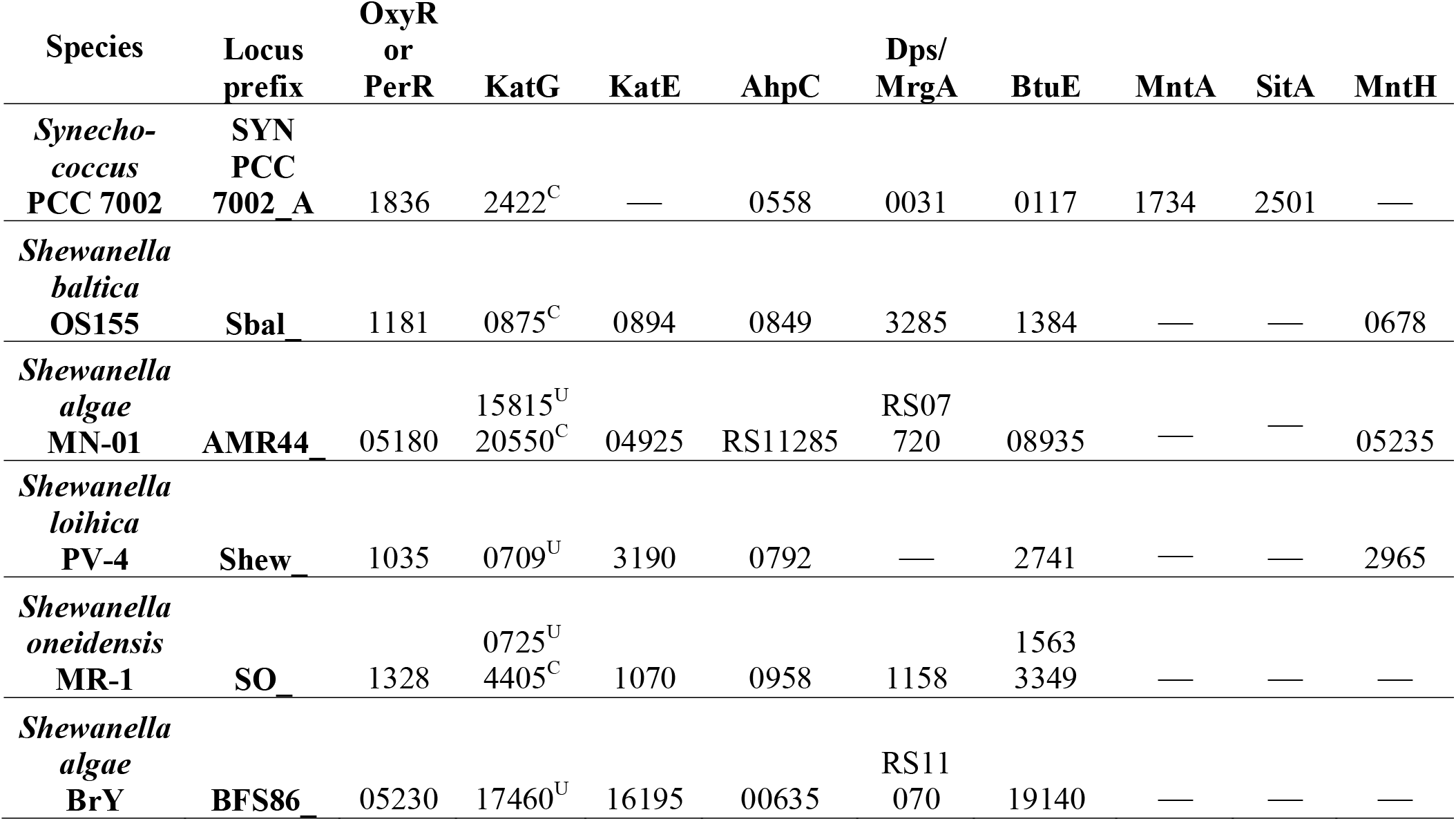
Locus tags of ROS response proteins in the bacterial species in this study. None contained Mn-catalase or Ni-superoxide dismutase. “—” indicates no homolog in genome. OxyR: hydrogen peroxide inducible gene activator; KatG: catalase-peroxidase (clade 1); KatE: periplasmic catalase (clade 3); AhpC: alkyl hydroperoxide reductase; Dps/MrgA: DNA-binding ferritin-like protein; BtuE: glutathione peroxidase; SitA and MntA: ABC-type Mn^2+^ transporters; MntH: NRAMP-type MntH Mn^2+^ transporter. KatG cellular localization based on PSORTb (Yu et al., 2010): C: cytoplasmic; U: unknown.

To identify genes in *Shewanella* that may have helped alleviate Fe^2+^ toxicity to *Synechococcus*, we compared the genomes of the *Shewanella* strains in our experiments. Notably, several *Shewanella* spp. contained catalases predicted to have multiple cellular locations, as previously observed for other microbial catalases (Hanaoka et al., 2013). Overall, the genomic inventory of catalase and peroxidase proteins was generally similar between the more protective and less protective species (**Table 1)**, suggesting (an)other mechanism(s) for ROS survival. We found 52 proteins in the best helpers (OS-155 and MN-01) that were not present in the other *Shewanella* strains, including genes for flagella, phenazine biosynthesis, and transporters **(Table S3)**. Flagella may be involved in the ROS-stress response in eukaryotes (Hajam et al., 2017), but their connection to ROS protection in bacteria, if any, is unknown. Phenazines are known to produce oxidative stress (Imlay, 2013), and can also mediate extracellular redox transfers (Hernandez et al., 2004; Wang and Newman, 2008), but are unlikely to be responsible for the protective effect because *Synechococcus* PCC 7002 also possesses the *phzF* gene for phenazine synthesis.

The high H_2_O_2_ sensitivity of *S. oneidensis* MR-1, which contains a similar repertoire of H_2_O_2_-scavenging enzymes as less H_2_O_2_-sensitive *Shewanella* spp., is thought to be due to its inability to actively transport and accumulate intracellular Mn^2+^ (Daly et al., 2004; Jiang et al., 2014). We surmised that differences in ROS scavenging rates between *Shewanella* strains may be due to differences in acquisition of Mn^2+^. We found MntH Mn^2+^ transporters in the genomes of the three top *Shewanella* helpers: OS-155, MN-01, and PV-4 (**Table 1**). *Shewanella* BrY and MR-1 lacked characterized Mn^2+^ transporters. *Synechococcus* PCC 7002 contained genes for the ATP-binding cassette (ABC) family Mn^2+^ transporters MntABCD and SitABCD transporter instead of MntH.

## Discussion

The rise of O_2_ and ROS from oxygenic photosynthesis would have severely stressed strictly anaerobic microbes (Khademian and Imlay, 2020), resulting in what was perhaps Earth’s first mass extinction. Experiments demonstrating that catalase-negative cyanobacteria (*Synechococcus* PCC 7002) grew poorly in >100 μM Fe^2+^ led to the idea that Fe^+^-rich oceans would have slowed cyanobacterial colonization of the ocean surface and possibly delayed global oxygenation (Swanner et al., 2015a). Our study confirms the previous finding that *Synechococcus* PCC 7002, originally isolated from marine mud, has impaired growth when Fe^2+^ was 180 µM and higher. We show that this Fe^2+^ toxicity can be alleviated by some strains of “helper” *Shewanella* spp., with the best protection afforded by *Shewanella* strains possessing the most varied sets of ROS-defense pathways (e.g. catalases, MntH transporters) and the highest rates of H_2_O_2_ degradation. Likely, this protection was afforded by *Shewanella* scavenging H_2_O_2_ prior to its reaction with Fe^2+^, thereby decreasing the production of damaging hydroxyl radicals.

Thus, our findings align with previous findings (Brown et al., 2010; Ward et al., 2019) that cyanobacterial colonization of early oceans would not have been hampered by micromolar Fe^2+^ concentrations, if Mn^2+^-transporting and H_2_O_2_-scavenging genes were present within the microbial communal gene pool. Early marine cyanobacteria, like modern terrestrial cyanobacteria, likely had myriad protections against Fe^2+^ and/or may have benefitted from the presence of co-existing “helper” bacteria to cope with the harmful byproducts produced by their own metabolism in a ferruginous ocean, which would have later been lost due to genome streamlining in marine cyanobacteria.

### Catalase-based protection

The ubiquity of catalase genes in the genomes of all the *Shewanella* strains we studied suggests that catalase accounts for the background protection provided by all *Shewanella* spp. tested. The enhanced protection provided by *Shewanella* spp. with similar catalase inventories implies that a mechanism other than catalase was likely at play, presumably at the level of gene expression. This process may also be related to the centralized regulation of H_2_O_2_-related genes in *Shewanella* spp. In *S. oneidensis* MR-1, the transcriptional regulator OxyR is key for suppression of Fenton chemistry by derepression of KatE and Dps (Jiang et al., 2014; Wan et al., 2018) whereas H_2_O_2_-based regulation is performed by multiple regulators (including PerR) in *Synechococcus* PCC 7002 and other cyanobacteria (Latifi et al., 2009).

A protective effect of proteobacterial catalase has previously been observed for the marine cyanobacterium *Prochlorococcus*, which grows in symbiosis with the gammaproteobacterium *Alteromonas* (Biller et al., 2016; Morris et al., 2011; Morris et al., 2008). (For more examples of microbe-microbe H_2_O_2_ protection, see review by Zinser (2018a).) Yet, unlike those long-lived catalase-based symbioses, the presence of *Shewanella* in our co-cultures was ultimately transient; *Synechococcus* gained the fitness advantage of protection from Fe^2+^ toxicity at the expense of *Shewanella*, whose population was eliminated from the system as *Synechococcus* grew. Indeed, previous attempts to co-culture *Shewanella* with cyanobacteria with ∼15 μM Fe^2+^, resulted in cyanobacterial dominance, with *Shewanella*’s growth yield was compromised by the presence of *Synechococcus* sp. 7002 even in the presence of organic carbon (Beliaev et al., 2014).

Thus, our co-culture experiments illustrate that cyanobacteria can benefit from the presence of “helper” proteobacteria under ferruginous conditions. This protection may have been one of the ways that cyanobacteria were able to cope with the harmful byproducts produced by their own metabolism as they incipiently colonized a ferruginous ocean, which would have no longer been necessary once cyanobacteria increased in numbers and seawater Fe^2+^ concentrations dropped. The precise levels of dissolved O_2_ prevailing on different spatial scales in the surface ocean prior to the GOE are not fully known. However, there is theoretical evidence to suggest that dissolved O_2_ would have been locally more than sufficient to support aerobic bacterial respiration (Olson et al., 2013; Reinhard et al., 2013a).

### Manganese to the rescue

One of the genes regulated by OxyR is the Mn^2+^ transporter MntH, which is used to accumulate intracellular manganese (Mn^2+^) as a potent ROS detoxification method (Anjem et al., 2009; Chen et al., 2008; Kehres et al., 2002). Unlike Fe^2+^, Mn^2+^ does not undergo Fenton-type reactions. Instead, Mn^2+^ has strong antioxidant properties (Cheton and Archibald, 1988) and is highly effective at protecting against H_2_O_2_-induced oxidative stress through multiple mechanisms (Aguirre and Culotta, 2012; Hansel, 2017; Horsburgh et al., 2002; Latour, 2015; Papp-Wallace and Maguire, 2006). Mn^2+^-carbonate and Mn^2+^-phosphate complexes can chemically disproportionate H_2_O_2_ (Archibald and Fridovich, 1982; Barnese et al., 2012; Stadtman et al., 1990). Mn^2+^-containing catalase, a very ancient member of the ferritin superfamily, detoxifies H_2_O_2_ (Klotz and Loewen, 2003; Zamocky et al., 2008). Under H_2_O_2_ stress, OxyR facilitates Mn^2+^ replacement of Fe^2+^ in ROS-sensitive enzymes, preventing their inactivation by Fenton chemistry (Anjem et al., 2009; Smethurst et al., 2020; Sobota and Imlay, 2011).

In monocultures, the rescued growth of *Synechococcus* PCC 7002 in two out of four of high Fe^2+^ treatments with 1 mM Mn^2+^ was likely related to the antioxidant properties of Mn^2+^, although the details of the protective mechanism, chemical or enzymatic, await further study. MntH transporters were found in the most protective *Shewanella* strains, but not in *Synechococcus* PCC 7002 (which instead encodes two ABC-type Mn^2+^ transporters) nor in the less protective *Shewanella* spp. **(Table S1)**. Although further experimentation at more environmentally relevant Mn^2+^ concentrations is needed, our initial findings generally support the hypothesis that elevated seawater Mn^2+^ in early Earth environments (∼5-120 µM; Holland, 1984; Johnson et al., 2016; Komiya et al., 2008; Liu et al., 2020) played a role in protecting marine cyanobacteria from ROS (Fischer et al., 2016; Lingappa et al., 2019).

### Modern microbial models for ancient physiologies

The choice of a model cyanobacterium for physiological experiments applicable to the Precambrian oceans is of great importance (Hamilton, 2019; Hamilton et al., 2016). Many terrestrial cyanobacteria thrive under the 10-100 µM Fe^2+^ concentrations predicted for Archean oceans (Brown et al., 2005; Ionescu et al., 2014; Thompson et al., 2019; Ward et al., 2019; Ward et al., 2017) and either possess multiple catalases **(Table S1)** and/or have novel defense mechanisms such as intracellular iron precipitation (Brown et al., 2010). In contrast, modern marine cyanobacteria (e.g. *Prochlorococcus*) tend to be genetically streamlined for specific modern oceanographic provinces (Partensky and Garczarek, 2010), which are extremely Fe^2+^-poor compared to modern terrestrial and ancient ecosystems.

The closest modern descendants of the ancestral cyanobacteria that evolved into modern marine plankton cyanobacteria are filamentous non-heterocystous *Synechococcales* (Sánchez-Baracaldo, 2015; Sánchez□Baracaldo and Cardona, 2020). KatG was likely present in ancestors of marine cyanobacteria (Bernroitner et al., 2009; Zamocky et al., 2012), whereas KatE was likely horizontally transferred from Proteobacteria and Planctomycetes to some cyanobacterial linages (e.g. Nostocales; Zamocky et al., 2012). *Synechococcus* PCC 7002 lacks Mn^2+^-catalase, which is widespread in terrestrial cyanobacteria (Ballal et al., 2020; Banerjee et al., 2012; Bihani et al., 2016; Chakravarty et al., 2016; Chen et al., 2020) **(Table S1)** and was likely present in early cyanobacterial lineages (Klotz and Loewen, 2003; Zamocky et al., 2012).

Previous genetic studies of Fe^2+^-induced oxidative stress have studied cyanobacteria that cannot cope with high Fe^2+^ and H_2_O_2_, e.g. *Synechocystis* PCC 6803 (Li et al., 2004; Shcolnick et al., 2009) in monoculture. In nature, ROS and O_2_ cycling are communal processes. Thus, models that include shared mechanisms of survival are important to consider on the early Earth, particularly as gene pools were more limited and were in the process of expansion. We advocate for future studies on more deeply branching cyanobacterial species with additional ROS defense mechanisms and on the molecular evolution of the Mn transporters and catalases discussed herein. We also encourage more explicit incorporation of microbial interactions in large-scale models of biogeochemical cycling on the ancient Earth.

## Materials and Methods

### Bacterial strains

*Synechococcus* sp. PCC 7002 was ordered from the Pasteur Culture collection of Cyanobacteria. *Shewanella oneidensis* MR-1 and *Shewanella algae* BrY were kind gifts from the lab of Dr. Thomas DiChristina (Georgia Institute of Technology). *Shewanella loihica* PV-4 was a kind gift from Dr. Jeffrey Gralnick (University of Minnesota).

### Experimental setup and growth conditions

*Synechococcus* sp. PCC 7002 was grown in serum bottles containing modified A+ medium (Stevens Jr. et al., 1973) with 10 g L^-1^ NaCl, TRIS buffer (pH 7.2), and 10 mM NH_4_^+^ as the nitrogen source. *Shewanella* spp. were grown overnight in lysogeny broth (LB, 10 g L^-1^ NaCl, 10 g L^-1^ tryptone, 5 g L^-1^ yeast extract) and transferred into serum bottles containing modified A+ medium with amino acids (20 mg L^-1^ L-serine, 20 mg L^-1^ L-arginine, and 20 mg L^-1^ L-glutamic acid), 20 mM lactate as electron donor, and 20 mM fumarate as electron acceptor. Bottles were flushed with 90% N_2_/10% CO_2_ and opened inside an anoxic chamber (5% CO_2_/4% H_2_/91% N_2_). Cultures were washed with anoxic A+ medium and combined at optical density of 600 nm (OD_600_) = 0.01. Co-cultures were grown in triplicate in 10 mL-well tissue culture plates inside the anoxic chamber (5% CO_2_/4% H_2_/91% N_2_). Cultures were mixed daily by gentle pipetting ∼50% of the volume three times in order to resuspend cells and particulate Fe(III) oxides; if not mixed regularly, PCC 7002 would grow at the bottom of the well. Light was provided with a fluorescent light in a 12:12 light:dark timer-controlled cycle. FeCl_2_ was added at a final concentration of 25, 500, or 1000 μM.

### Cyanobacterial quantification by flow cytometry

Cell numbers of *Synechococcus* sp. PCC 7002 were quantified in an LSR Fortessa flow cytometer using FACSDiva™ (BD Biosciences, CA). At each time point, 200 μL of culture was loaded into a 96-well plate inside the anoxic chamber, covered in parafilm to minimize Fe^2+^ oxidation, transported to the cytometer, and mixed twice in the cytometer. Samples (10 μL) were injected and run at a rate of 0.5 μL s^-1^. Cyanobacteria were detected by phycocyanin/chlorophyll autofluorescence using blue (488 nm) and yellow-green lasers (561 nm) measured at 655-684 nm (Hill et al., 2017). Optimization and calibration of the quantification parameters were achieved using yellow-green 1 μm microspheres (441/485 ex/em; Polysciences, PA). Live cyanobacteria were also quantified using Syto9 using FITC filters. Events above the thresholds of PerCP and FITC were considered live cyanobacteria. Propidium iodide could not be used to identify “dead’ cyanobacteria, as the emission spectra overlapped with that of their autofluorescence. Due to spectral overlap with iron particles, *Shewanella* cells could not be accurately quantified by cytometry.

### H_2_O_2_ resistance assays

Six *Shewanella* strains were incubated in 96-well plates with minimal M1 media (Myers and Nealson, 1988) with lactate (10 mM) or acetate (10 mM) as electron donor under oxic conditions. Growth was monitored periodically (every 1-2 hours) by OD_600_ in a spectrophotometer with plate-reading capacity (Tecan, Switzerland). Hydrogen peroxide was added after an initial period of growth for 4 hours, at a final concentration of 10, 25, 50, 100, or 250 μM, after which growth continued to be monitored. (Note: these concentrations are 10-10,000x higher than most natural waters, which rarely exceed 1 µM H_2_O_2_ (Cooper et al., 1988)).

### H_2_O_2_ scavenging assays

We compared the abilities of the *Shewanella* spp. to remove H_2_O_2_ from their environment in cell suspensions. Strains were seeded in lysogeny broth (LB, 10 g L^-1^ NaCl, 10 g L^-1^ tryptone, 5 g L^-1^ yeast extract, Sigma-Aldrich) at 30°C with shaking overnight, harvested by centrifugation at 12,300 x g, washed, and transferred into minimal M1 medium amended with 20 mM lactate at OD_600_ = 0.02. Cells were incubated at 30°C with shaking until harvesting at mid-log phase (OD_600_ = 0.15-0.35), washed twice with minimal medium, then inoculated at OD_600_ = 0.05 into a 24-well plate holding 2 mL minimal M1 medium amended with 20 mM lactate and various concentrations of H_2_O_2_ (0-5000 μM). Samples were collected every 3-5 minutes and analyzed immediately for exogenous H_2_O_2_ using the resorufin-horseradish peroxidase colorimetric assay (Zhou et al., 1997). Plates were incubated under oxic conditions at room temperature with shaking for the duration of the experiment (30-200 min). H_2_O_2_ disappearance followed an exponential decay **(Eq. 2)**. First-order apparent rate constants (k) were obtained by plotting the data as shown in **Eq. 3**, where k is the slope of the graph with ln[H_2_O_2(t=0)_/H_2_O_2(t=n)_] on the y-axis and time on the x-axis.

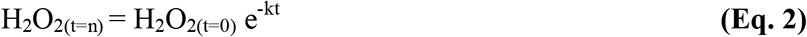

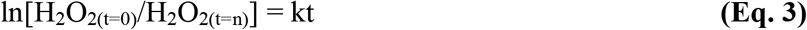

### Unique proteins

Proteins present in *Shewanella algae* MN-01 and *Shewanella baltica* OS155 and absent in *Shewanella algae* BrY, *Shewanella oneidensis* MR-1, and *Shewanella loihica* PV-4 were identified using the Protein Families tool in PATRIC using three protein family databases: PATRIC cross-genus families (PGfams), PATRIC genus-specific families (PLfams), and FIGfam.

### Synechococcus monoculture experiments

To determine the influence of 1 mM Mn^2+^ on *Synechococcus* PCC 7002 growth with and without 1 mM Fe^2+^, *Synechococcus* was grown in modified A+ medium containing 82 mM bicarbonate in Hungate tubes with bromobutyl rubber stoppers containing 95% N_2_/5% H_2_ headspace with constant shaking at 200 rpm under constant light. Growth was determined by measurement of optical density at 750 nm (OD_750_).

## Supplemental tables

**Table S1. Cyanobacterial genomes containing MnKat, KatE, or KatG**. Appended as separate spreadsheet.

**Table S2. Proteins that are present in *Shewanella algae* MN-01 and *Shewanella baltica* OS155 and are not present in *Shewanella oneidensis* MR-1, *Shewanella algae* BrY, and *Shewanella loihica* PV-4**. Appended as separate spreadsheet.

## Acknowledgements

This research was funded by NASA Exobiology grant NNX14AJ87G, NASA Astrobiology Institute grant 13-13NAI7_2-0027, and a NASA Astrobiology Postdoctoral Fellowship to NS. We thank Jennifer Thweatt for advice on culturing *Synechococcus* sp. PCC 7002. We thank Amit Reddi for allowing us to use his plate reading spectrometer. We thank Sommer Durham for technical assistance with the cytometer. We thank Patricia Sanchez-Baracaldo and Manuel Kleiner for helpful discussions.

